# Peri-adolescent THC exposure does not lead to anxiety-like behavior in adult mice

**DOI:** 10.1101/2020.08.31.274274

**Authors:** Matija Sestan-Pesa, Marya Shanabrough, Tamas L. Horvath, Maria Consolata Miletta

## Abstract

As marijuana use during adolescence has been increasing, the need to understand the effects of its long-term use becomes crucial. Previous research suggested that marijuana consumption during adolescence increases the risk of developing mental illness, such as schizophrenia, depression, and anxiety. Ghrelin is a peptide produced primarily in the gut and is important for feeding behavior. Recent studies have shown that ghrelin and its receptor, the growth hormone secretagogue receptor (GHSR) play important roles in mediating stress, as well as anxiety and, depression-like behaviors in animal models. Here, we investigated the effects of chronic Tetrahydrocannabinol (THC) administration during adolescence (P42-55), in GHSR (GHSR^-/-^) knockout mice and their wild type littermates in relation to anxiety-like behaviors. We found that continuous THC exposure during peri-adolescence did not lead to any significant alterations in anxiety-like behavior of male adult mice, regardless of genotype. These data indicate that in the presence of intact GHSR signaling, THC exposure during peri-adolescence has limited if any long term impact on anxiety-like behaviors in mice.

## 1. Introduction

Adolescence is the developmental period of transition between childhood and adulthood, on average starting at age 12 and ending at age 17 [1, 2]. This period is marked with significant neuroplasticity and changes happening, primarily in the prefrontal cortex and limbic regions, which contribute to subsequent adult behavior and cognitive functions [1, 3]. Cannabis use among adolescents is very high, with 9.4% of 8^th^ graders, 23.9% of 10^th^ graders, and 36.5% of 12^th^ graders reporting cannabis use in the last 12 months in 2016 [4]. This event is concerning as cannabis use in adolescence has been associated with several negative outcomes in adulthood, particularly as a risk factor for developing cognitive impairment and mental illness later in life [5]. Anxiety appears to be the most common complication arising from heavy cannabis use, with up to 20% of cannabis users having anxiety [6]. This is higher than the prevalence of anxiety in the general population, which is estimated to be around 6-17%. There is also a positive significant association of anxiety with cannabis consumption and misuse [7]. Cannabis use during adolescence has also been shown to double the risk of developing anxiety-related symptoms later in life, especially if cannabis use started before the age of 15. Girls appear to be more likely to develop these symptoms, indicating a gender difference in the long-term effects of cannabis [8, 9].

In animal models, the connection between cannabis use and anxiety is not as consistent as in humans. In some cases, [10]. Chronic administration of CB1R agonists (e.g., Tetrahydrocannabinol (THC), CP-55,940, and WIN55, 212-2) in adult rats decreased anxiety-like behavior [11-13]. On the other hand, animals treated with CB1R agonists (CP-55,940 and THC) only several days earlier than the above-mentioned animals, showed an increase in anxiety-like behaviors in adulthood [14].

The endocannabinoid system, which consists of cannabinoid receptors (CB1R, CB2R) and their naturally occurring ligands (anandamide and 2-arachidonoyl glycerol), appears to play a modulating role on proliferation and differentiation of progenitor cells, neuronal migration, axonal guidance, fasciculation, positioning of cortical interneurons, neurite outgrowth and morphogenesis [15, 16]. During development for infancy to young adulthood, CB1R expression increases dramatically in regions such as the prefrontal cortex, striatum, and hippocampus [17]. Imaging studies have shown decreased cortical thickness in the right superior PFC, bilateral insula and bilateral superior cortices in adolescent cannabis users compared to adolescents who do not use cannabis [18], as well as a decrease in volume of the right medial orbitofrontal cortex [19] and bilateral hippocampus [20, 21].

Ghrelin’s role in regulating mood appears to be a very complex one and there is reason to believe it has a dual role in regulating anxiety. In some cases, injecting ghrelin centrally increased anxiety-like behavior assessed by elevated plus maze [22], while other reports suggest the opposite effect, with ghrelin injections showing a decrease of anxiety-like behavior as assessed by elevated plus maze [23]. This discrepancy might be related to the timing of the behavioral experiments. Another factor that contributes to modulate ghrelin’s effect on behavior is food availability, with ghrelin increasing locomotion in the absence of food [24] and decreasing locomotion in the presence of food [25]. Findings in ghrelin knockout mice also demonstrate this ambiguous relationship between ghrelin and anxiety. Ghrelin knockout (Ghr -/-) mice appear to be less anxious than their wild-type counterparts under non-stressed conditions but displayed more anxious behavior under mild stress conditions (15-min restraint) [26]. Of note is that stress increases ghrelin and corticosterone concurrently. GHSR and ghrelin knockout mice showed decreased plasma levels of corticosterone after chronic social defeat stress and acute restraint stress, as well as increased anxiety-like behavior [26, 27]. Taken together these findings suggest that ghrelin and GHSR are important for the ability of animals to cope with anxiety-inducing stressors.

Ghrelin also has neuroprotective properties, as it increases hippocampal neurogenesis [28], and GHSR -/- mice have reduced cell proliferation and survival in the dentate gyrus of the hippocampus, as well as an increase in apoptosis [29]. This might be related to its role in modulating anxiety. Since cannabis consumption has been correlated to changes in the hippocampus [17, 20, 21], we decided there was sufficient neurobiological rationale to warrant an investigation into how GHSR -/- mice (and their wild type counterparts) would respond to chronic THC administration during adolescence. To investigate the long-term effects of THC on behavior relating to anxiety, we exposed the animals to 10mg of THC daily (via pulmonary route) during sexual maturation (6-8 weeks), which roughly corresponds to adolescence in humans. After 14 days of THC administration, animals could recover for an additional 4 weeks. At 12 weeks of age, behavioral testing was performed to evaluate any long-term effects from THC administration (Fig. 1a). Male and female mice were used to evaluate the possibility of males and females being affected differently since the literature suggests that females may be more vulnerable to THC’s effect on anxiety [8, 9].

**Figure 1.**
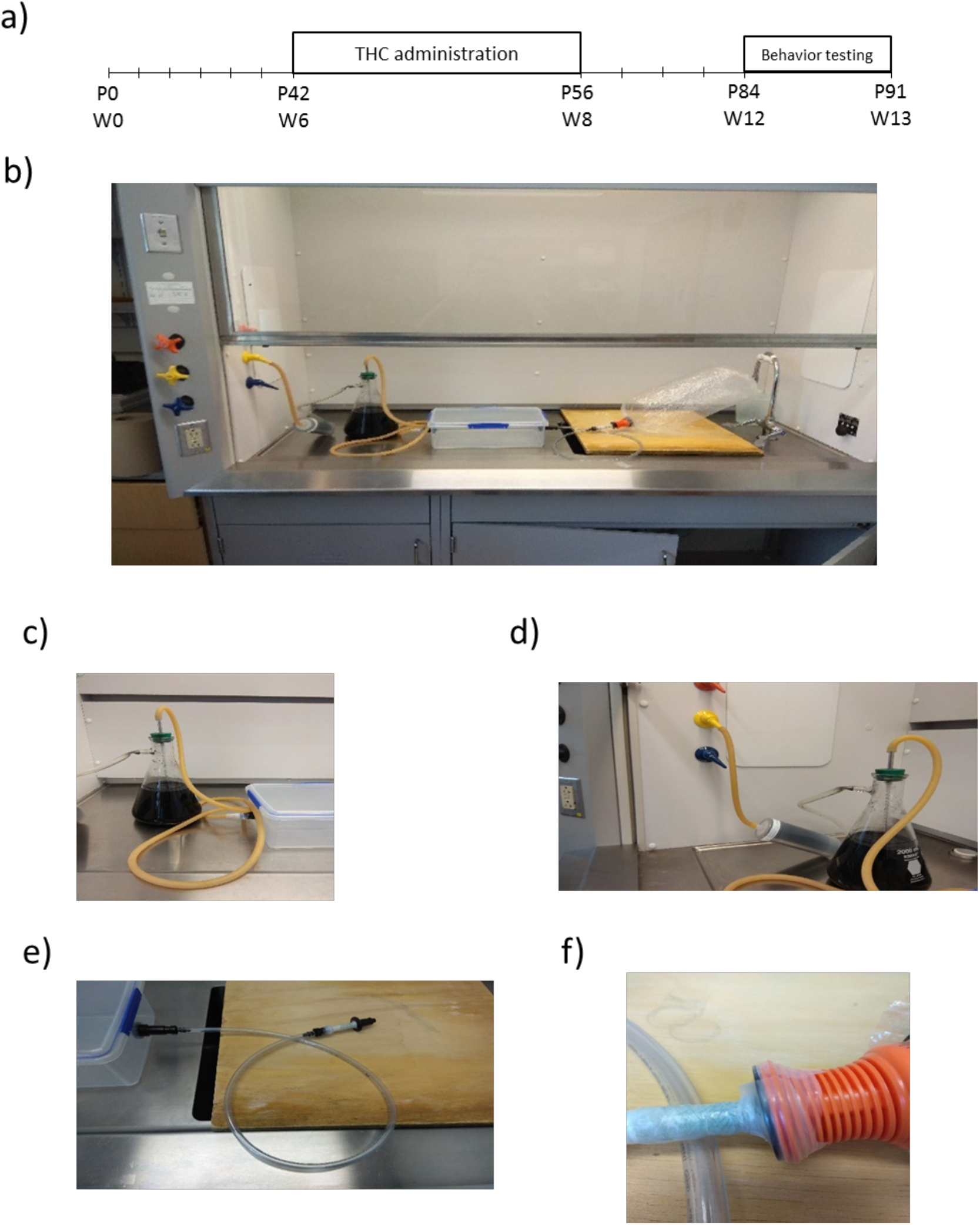
Experiment design and equipment for THC administration. **a)** Experimental design, the time course for THC (or vehicle) administration and behavioral testing **b)** Entire apparatus used to administer THC and vehicle under a chemical hood **c)** The tubing leading from administration box to activated charcoal trap **d)** The tubing leading from activated charcoal trap to the activated charcoal filter, which then leads to the vacuum line **e)** Open-ended tube with Volcano mouthpiece attached **f)** The balloon attached to the open-ended mouthpiece, sealed with parafilm

## 2. Materials and Methods

### 2.1. Materials

To closely mimic human THC consumption, we used a formulation of THC (3.62% THC, 6.47% THCA, a total of 101 mg/ml of THC) with a minimal content of terpenes (β-myrcene 0.06%, β-caryophyllene 0.64%, humulene 0.39%; a total of 1.09% terpenes) dissolved in PEG 400, designed for use with a commercially available vaporization apparatus. PEG 400 with terpenes was used as the vehicle for the control group. Connecticut Pharmaceutical Solutions, LLC, Portland, Connecticut (a state-licensed grower) provided the compounds through the Connecticut Medical Marijuana Research Program.

### 2.2. Animals

The Institutional Animal Care and Use Committee of Yale University approved all experiments. Mice were kept under standard laboratory conditions with free access to standard chow food and water except during behavioral testing. GHSR^-/-^ mice were previously generated in our lab [30]. THC was administered to animals from 6 to 8 weeks of age and behavioral testing was performed at 12 to 13 weeks of age (Fig.1a). Animals were placed into 2 treatment groups (vehicle and 10mg THC) for each genotype (wild-type and knockout).

### 2.3. THC administration

Most adolescents smoke cannabis, therefore we decided to mimic smoking as a method of administration of THC. We used commercially available vaporization equipment for marijuana administration, commonly used by marijuana consumers. Previously described experiments used the Volcano® Vapourization device (Storz and Bickel, GmbH and Co., Tuttlingen, Germany) to administer ethanol dissolved THC to lab animals [31, 32]. All administrations of THC and vehicle were done under a chemical hood to prevent cross-contamination (**Fig. 1b**). Mice were placed, in groups of 2-4, inside a closed chamber (33cm x 20.3cm x 10.2cm) with valves and tubing on two of the narrower sides. To further minimize vapor escaping, on one side the tubing leads to an improvised activated charcoal trap, leading to an activated charcoal filter, leading to the vacuum line (**Fig. 1c, 1d**). On the other side, tubing was open-ended with a Volcano Vaporizer mouthpiece fixed to it (**Fig. 1e)**. The mouthpiece was used to release the seal on the balloons, which were filled with a vapor containing THC (or vehicle; note that the content of the balloons is mostly air so that the animal was always normoxic while in the chamber). Parafilm was used to seal the connection making it airtight (**Fig. 1f**). Animals were exposed to the vapor for 5 min, with half of the balloon being emptied at the beginning and the other half being emptied at 2:30 min of exposure. After the exposure, animals were quickly removed from the chamber and the vacuum line was turned on to remove any residual vapor left. The inside of the box was cleaned with 70% ethanol between each group of animals.

### 2.4. Behavioral assessments

Open field and elevated zero maze were used to establish any behavioral phenotypes induced by THC administration. All testing was done in the light phase of the cycle from 1 to 7 pm.

#### 2.4.1. Open field

The open field apparatus was a square, polyurethane box (35.5cm x 35.5cm x 30cm). The animal was placed in the center of the apparatus. General locomotion parameters (distance traveled, locomotion speed, time mobile) and parameters relating to anxiety (freezing time; time spent, distance traveled, and entries into central and periphery zones) were recorded for 10 min. The apparatus was cleaned with 70% ethanol after each animal exposure. ANY-Maze software (Stoelting Company, Wood Dale, IL) was used to record and analyze the behavioral data.

#### 2.4.2. Elevated zero maze

The elevated zero mazes was an elevated (60cm high) ring-shaped runway (5cm wide), with 2 equally sized (25% of the runway length) sections closed off by walls (40cm high) opposite each other. The other two sections were open. The maze was equally illuminated in all four sections. Mice were placed on the center of one of the open sections, facing one of the closed sections, and allowed to explore the maze for 5 min. The apparatus was cleaned with 70% ethanol after each animal exposure. ANY-Maze SoftwareTM (Stoelting Company, Wood Dale, IL) was used to record and analyze the behavioral data.

### 2.5. Statistics

Prism 8.0 was used to analyze data and plot figures. In our study, the sample sizes were chosen based on the common practice in animal behavior experiments[33]. Two-way ANOVA was used as the other tests unless stated otherwise. When significant, a multiple comparisons post-hoc test was used. Statistical data are provided in the figures. P < 0.05 was considered statistically significant.

## 3. Results

### 3.1. Open field

The open field exploration test represents a unique opportunity to systematically assess novel environments exploration, general locomotor activity and provides initial screening for anxiety-related behavior in rodents[34]. Two factors influence anxiety-like behavior in the open field. The first is social isolation resulting from the physical separation from cage mates when performing the test. The second is the stress created by the brightly lit, unprotected, novel test environment[35, 36].

In our experimental setting, parameters that evaluate the general locomotion of the males showed no differences amongst the groups, suggesting locomotion was unaffected by THC exposure (**Fig. 2a, b, c, 2e**). The only exception was the time animals spent freezing inside the apparatus. Two-way ANOVA analysis showed a significant effect of THC treatment (*, p= 0.0244), but there were no post-hoc significant differences, making it difficult to deduce if this effect was due to variability in animal behavior, or an underlying trend of the relevant difference caused by THC exposure. (**Fig. 2d**) Preliminary results on female mice showed no differences as well. (**Fig. S1a-e**). Parameters in the open field indicative of anxious behavior showed no differences in the male mice, suggesting no changes in anxiety-like behavior (**Fig. 3a-e**). Preliminary results on female mice show no significant differences as well (**Fig. S2a-f**).

**Figure 2.**
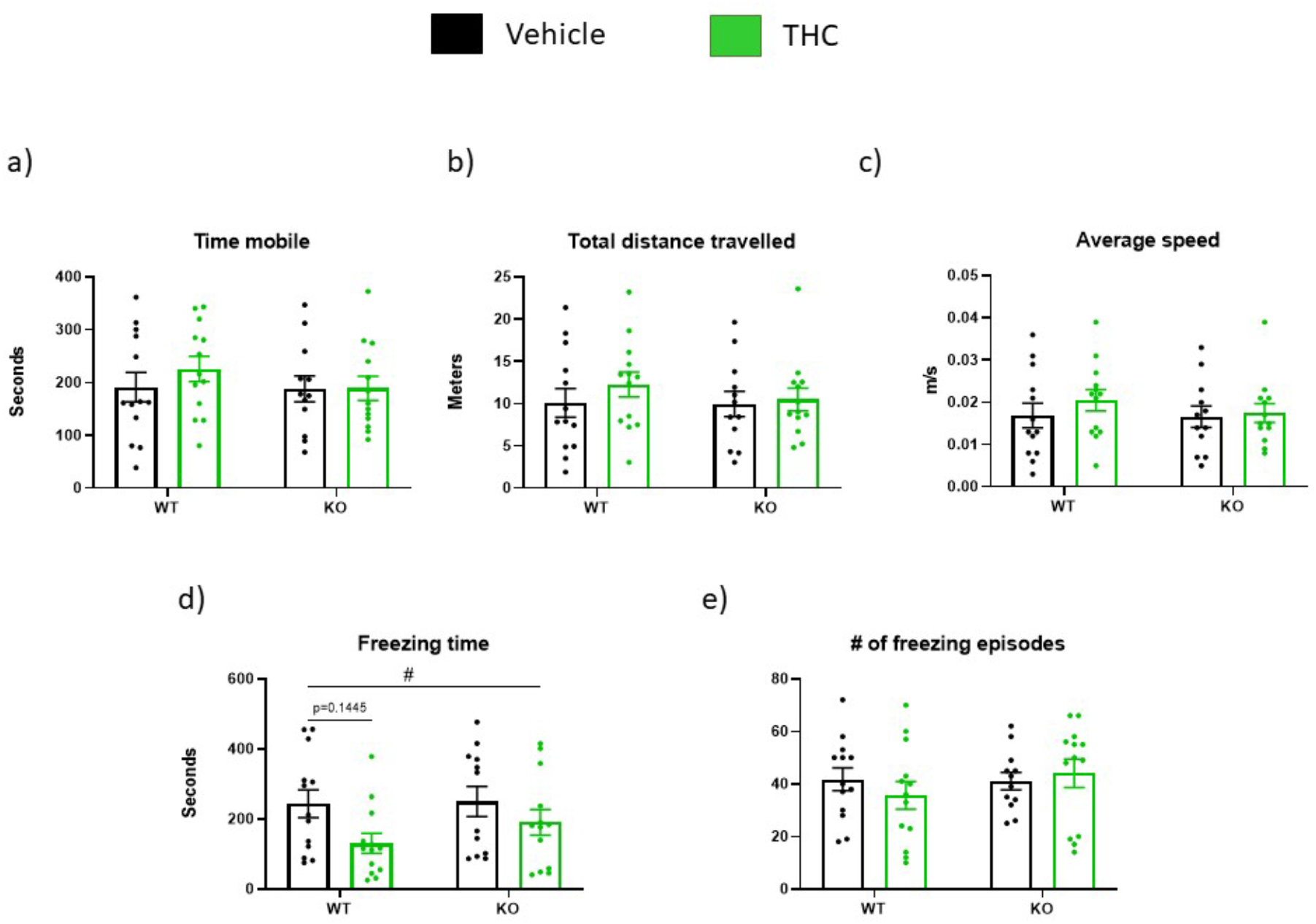
Open field, general locomotion. WT-vehicle n=13, WT-THC n=13, KO-vehicle n=12, KO-THC n=13; **a)** Time mobile **b)** Total distance traveled **c)** Average speed **d)** Freezing time **e)** Number of freezing episodes. Data are presented as mean values ± SEM. # p< 0.05 effect of THC treatment (two-way ANOVA analysis). Post-hoc comparison indicated by p values where appropriate.

**Figure 3.**
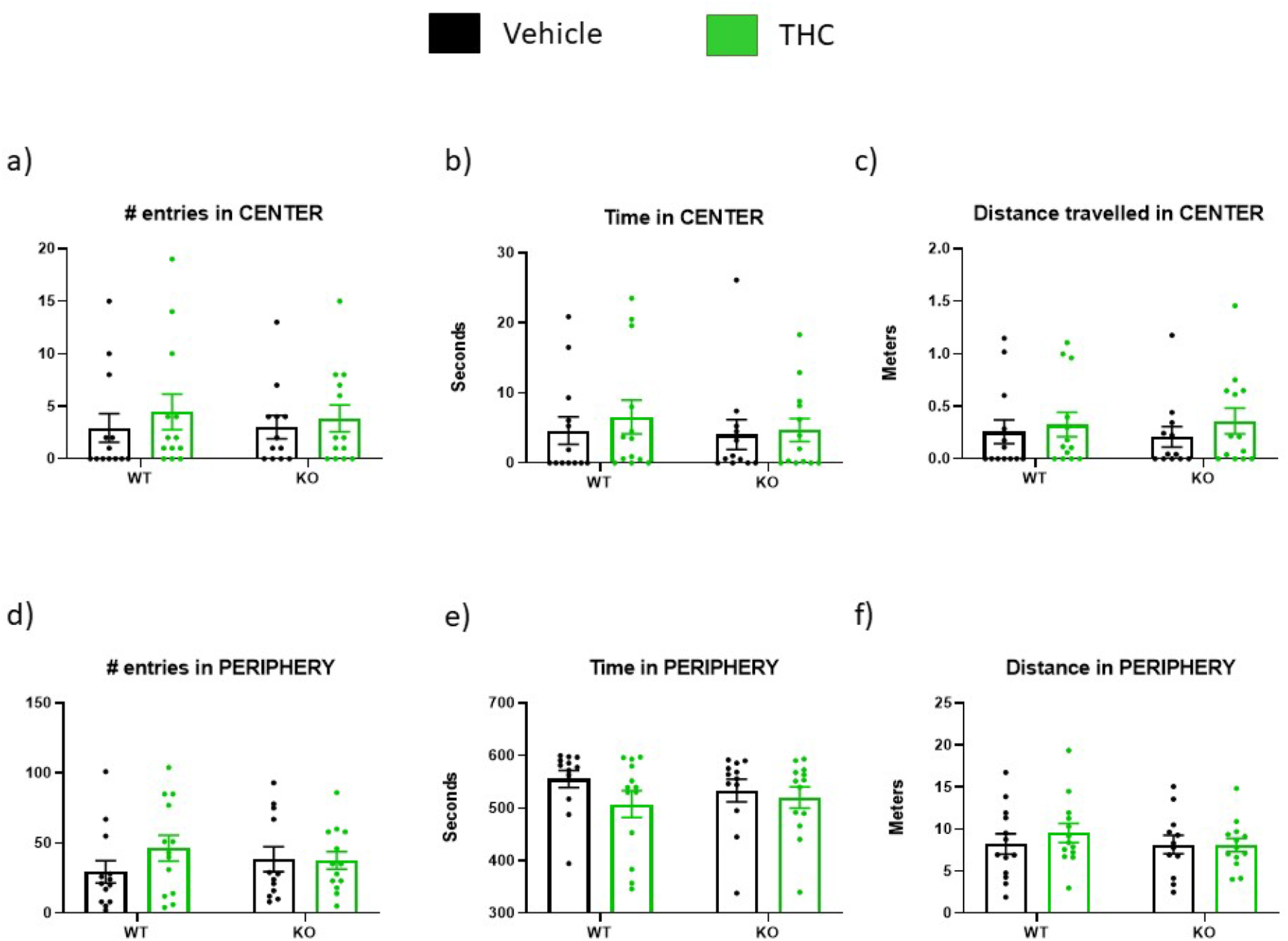
Open field, anxiety-related parameters. WT-vehicle n=13, WT-THC n=13, KO-vehicle n=12, KO-THC n=13; **a)** Number of entries in the central zone **b)** Time spent in central zone **c)** Total distance traveled in central zone **d)** Number of entries in peripheral zones **e)** Time spent in peripheral zones **f)** Total distance traveled in peripheral zones. Data are expressed as mean ± SEM *p< 0.05. Two-way ANOVA

### 3.2. Elevated Zero Maze

Elevate zero maze is the master test for assessing anxiety-like behavior in mice. The test exploits the natural tendencies of mice to explore novel environments[34, 37].

There were no significant effects in the male mice, suggesting THC exposure had no significant lasting effects on anxiety (**Fig. 4a-e**). Two-way ANOVA analysis showed that the effect of THC treatment was close to significance in the parameters of Time Spent in Open Arms and Distance Travelled in Open Arms (p= 0.0758 and p= 0.0746 respectively), however, no post-hoc difference was close to significance, suggesting this trend was probably caused from the natural variability of mouse behavior (**Fig. 4b, 4c**). Preliminary experiments on female mice indicate a possible trend, with two-way ANOVA showing significant effects of THC treatment in Time Spent in Open Arms (*, p=0.0151) and Time Freezing in Open Arms (*, p=0.0268) however; there were no significant post-hoc differences between individual groups. (**Fig S3B, S3E**) Other parameters show no differences or trends. **(Fig S3A, S3C, S3D)**

**Figure 4.**
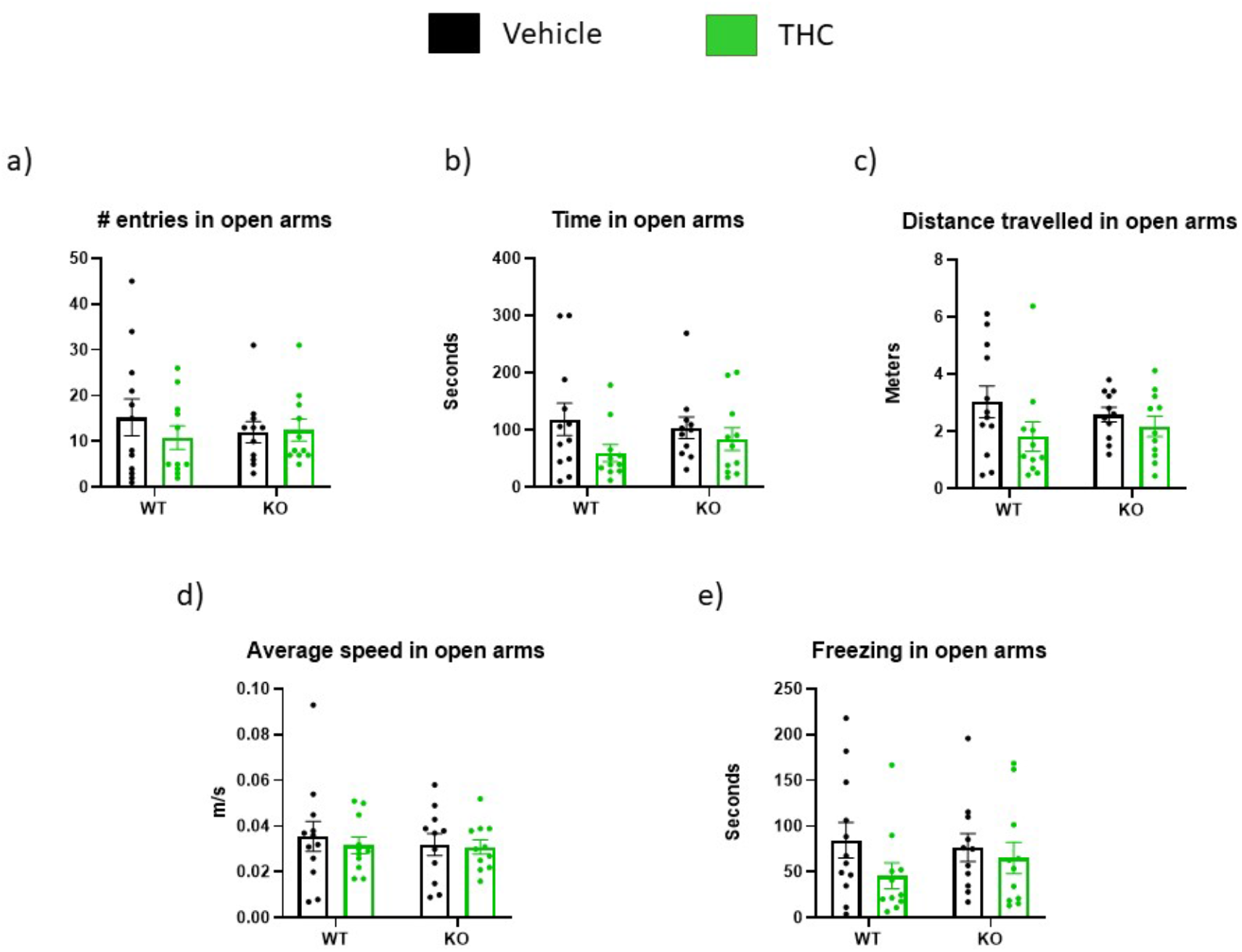
Elevated Zero Maze. WT-vehicle n=12, WT-THC 10 mg n=11, KO-vehicle n=11, KO-THC 10 mg n=11 **a)** Number of entries in open arms **b)** Time spent in open arms **c)** Distance traveled in open arms **d)** Average speed in open arms **e)** Time spent freezing in open arms. Data are expressed as mean ± SEM *p< 0.05. Two-way ANOVA.

## 4. Discussion

Upon examination of the data from the open field test, we can conclude that THC exposure during peri-adolescence did not affect the baseline locomotion of males, in adulthood. While there was an effect of THC treatment in the parameter of the time-spent freezing, no significant post-hoc differences were found suggesting this effect was likely due to the natural variability of animal behavior. Parameters indicative of exploratory behavior and anxiety showed no differences suggesting peri-adolescent THC exposure does not induce long-term changes in the anxiety of male mice.

The results from the elevated zero maze further corroborate a lack of any significant long-term effects of peri-adolescent THC exposure on exploratory and anxiety-like behavior in male adults. These results and those from the open field test, suggest that impairment of the ghrelin system does not confer an increased risk of developing THC induced anxiety in adult males.

Preliminary experiments on female mice showed similar results as the males, except concerning the elevated zero maze data, with THC treated animals spending more time on the open arms. These data would suggest a decrease in anxiety-like behavior due to peri-adolescent THC exposure. But there are a few reasons we should be skeptical of this interpretation. Firstly, these findings, although close to statistical significance, are still underpowered in terms of sample size, thus any conclusion made on this dataset would be misleading. Secondly, the open field data showed no signs of difference in exploratory and anxiety-like behavior. Finally, the second parameter of the elevated zero maze that showed a significant effect of treatment (time spent freezing in open arms), is suggestive of the opposite (more freezing would indicate higher levels of fear/anxiety). This would imply that more time spent on the open arms is an artifact caused by higher levels of fear leading the animals to freeze on the open arms, confounding the results. The other parameters, such as distance traveled, showed no signs of any similar trends. If this trend is reflective of a true difference, it may be related to ghrelin’s neuroprotective role in some parts of the brain (hippocampus, *substantia nigra*, hypothalamus, raphe nuclei), since the effect is only close to significance in the knockout mice [38, 39].

Since there is concern about the effects of THC consumption among adolescents, it would be important to verify the lack of significant long-term alterations of behavior (concerning impaired/intact GHSR signaling) induced by THC, as presented in this paper. Changing conditions such as the time of THC exposure, concomitant stress exposure and presence/lack of food could clarify if there are any relevant conditions under which THC can significantly alter long-term behavior. The time point of behavioral testing is also important since there may be significant alterations in behavior but at an earlier time point. By the time we tested these animals, they may have already recovered from any relevant impairments to their measurable behavior. To further address the possible sex differences and ghrelin’s role in them, we suggest to accompany the THC exposure with a variety of adjunct treatments, such as sex hormone inhibitors and ghrelin, as well as repeating this in ghrelin knockout mice to elucidate the mechanism of the sex difference and how ghrelin and its receptor may play a role in this.

It will also be important to investigate if any of these changes in behavior are accompanied by changes in areas of the brain involved in these behaviors, such as the hippocampus, amygdala, or the prefrontal cortex. Lastly, the tester should employ more behavioral tests such as pre-pulse inhibition, marble burying, and tail suspension to investigate whether THC exposure in peri-adolescence affects behaviors not directly relating to anxiety but more related to sensory gating, compulsiveness and mood regulation.

## Supplementary Materials

**Figure S1.**
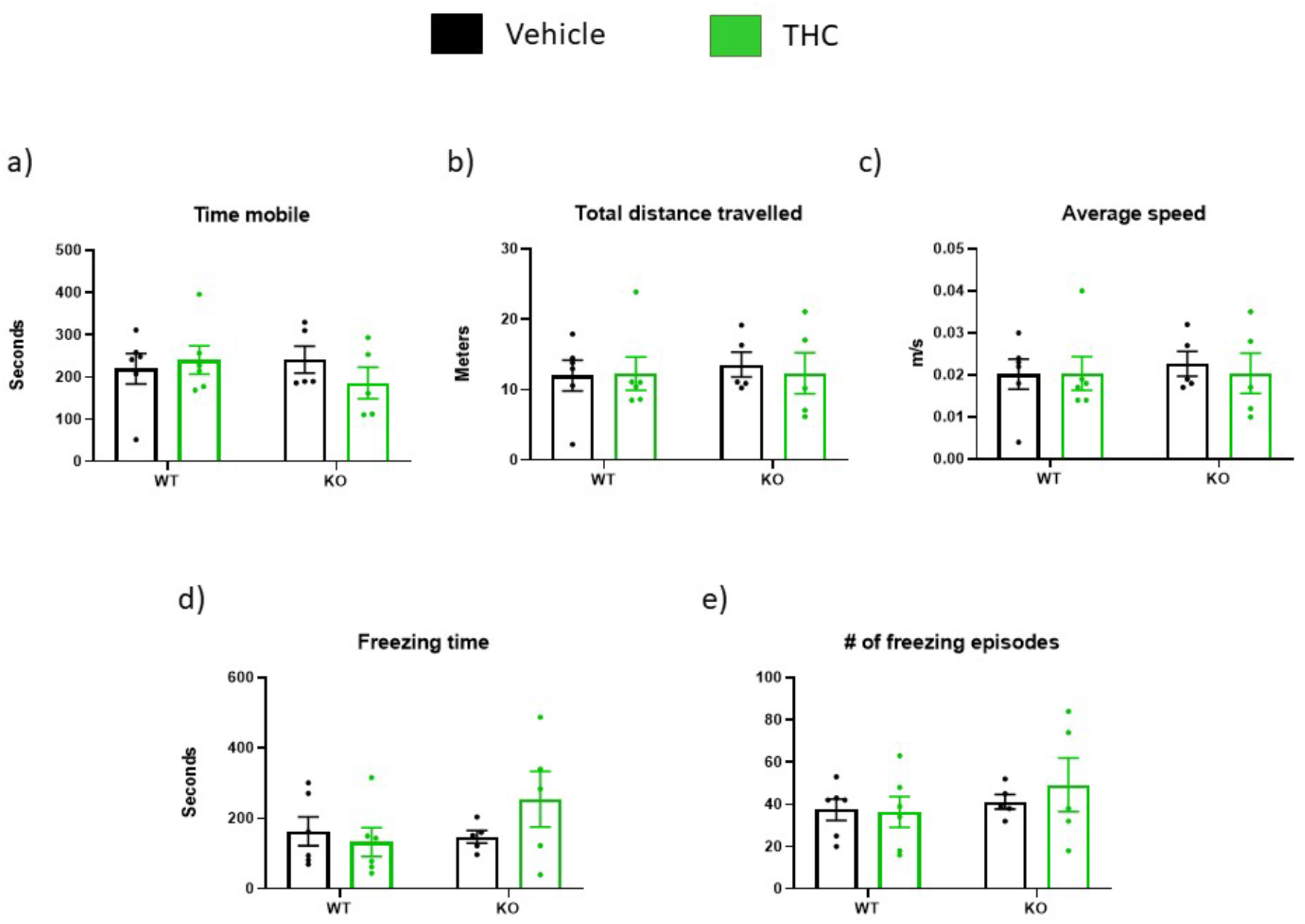
Females, open field, general locomotion. WT-vehicle n=6, WT-THC n=6, KO-vehicle n=5, KO-THC n=5; **a)** Time mobile **b)** Total distance traveled **c)** Average speed **d)** Freezing time **e)** Number of freezing episodes. Data are expressed as mean ± SEM *p< 0.05. Two-way ANOVA.

**Figure S2.**
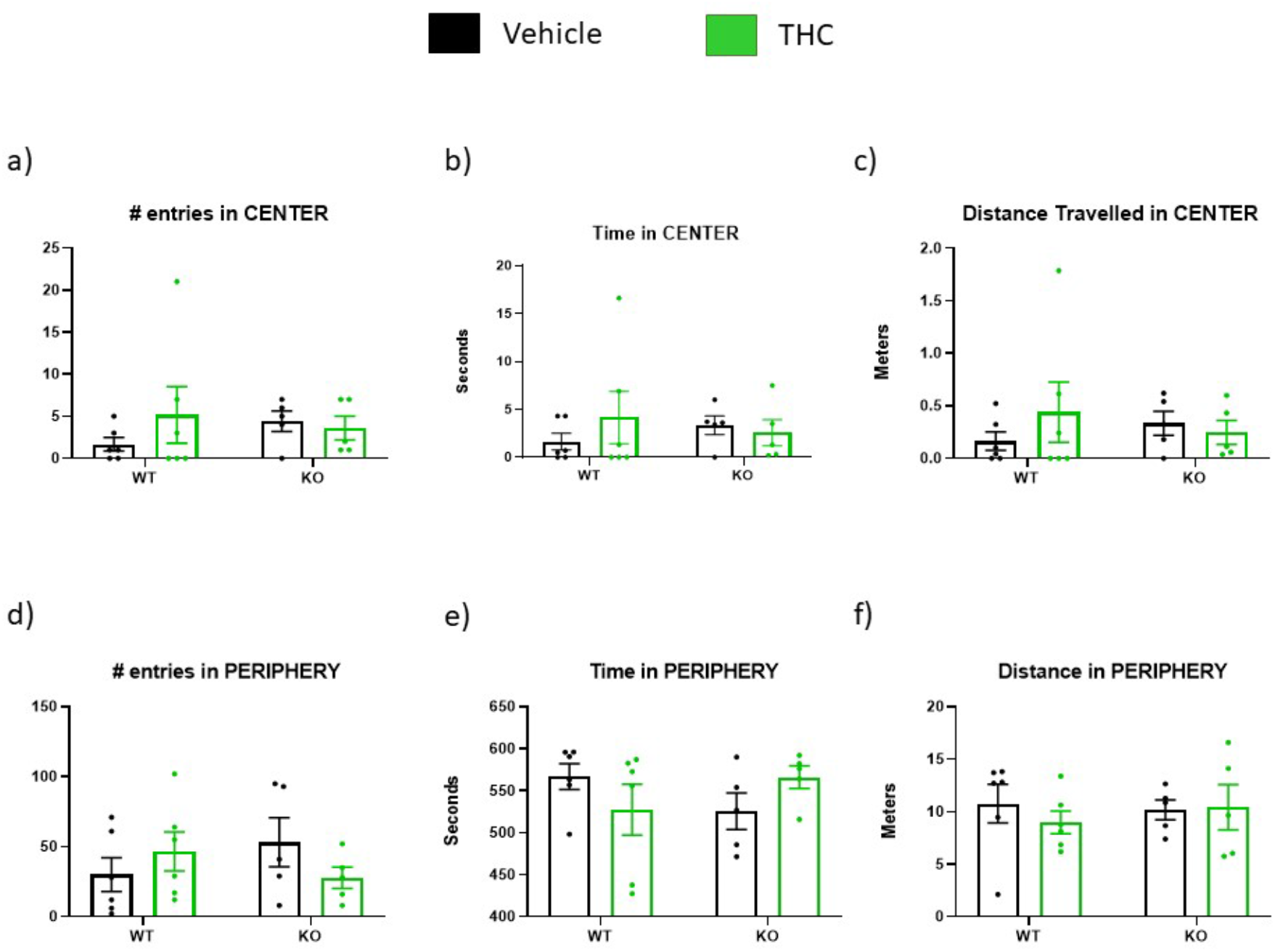
Females, open field, anxiety-related parameters. WT-vehicle n=6, WT-THC n=6, KO-vehicle n=5, KO-THC n=5; **a)** Number of entries in the central zone **b)** Time spent in central zone **c)** Total distance traveled in central zone **d)** Number of entries in peripheral zones **e)** Time spent in peripheral zones **f)** Total distance traveled in peripheral zones. Data are expressed as mean ± SEM *p< 0.05. Two-way ANOVA.

**Figure S3.**
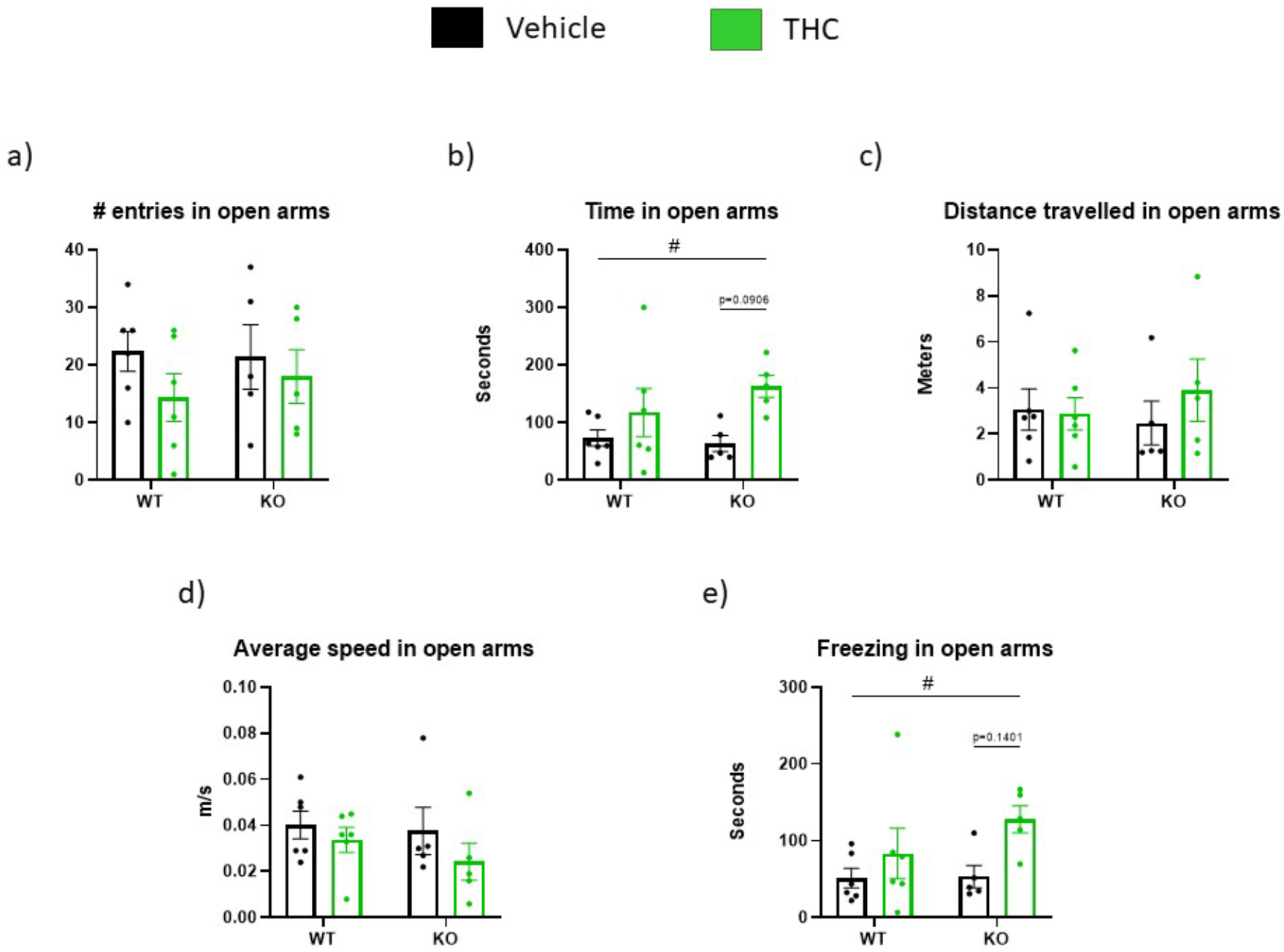
Females elevated zero maze. WT-vehicle n=6, WT-THC n=6, KO-vehicle n=5, KO-THC n=5 **a)** Number of entries in open arms **b)** Time spent in open arms **c)** Distance traveled in open arms **d)** Average speed in open arms **e)** Time spent freezing in open arms. Data are presented as mean values ± SEM. # p< 0.05 effect of THC treatment (two-way ANOVA analysis). Post-hoc comparison indicated by p values where appropriate.

## Author Contributions

T.L.H. developed the concept. M.C.M. M.S-P developed the experimental strategy with input from T.L.H. M. S. provided throughout technical support. M.C.M., and M.S-P conducted experiments and. analyzed data. M.S-P, M.C.M. and T.L.H. wrote the paper with input from all authors

## Acknowledgments

We would like to thank Connecticut Pharmaceutical Solutions, LLC for providing us with the compounds used in this study.

